# Neuropeptide-mediated temporal sensory filtering in a primordial nervous system

**DOI:** 10.1101/2024.12.17.628859

**Authors:** Livia S. Wyss, Samuel R. Bray, Bo Wang

**Author notes:** **Author contributions.** Conceptualization: LSW, SRB, and BW; Methodology: LSW, and SRB; Investigation and formal analysis: LSW; Writing: LSW, and BW, with feedback from all other authors; Funding acquisition: BW; Supervision: BW. **Competing Interest Statement:** The authors declare no competing interests.

## Abstract

Sensory filtering – prioritizing relevant stimuli while ignoring irrelevant ones – is crucial for animals to adapt and survive in complex environments. While this phenomenon has been primarily studied in organisms with complex nervous systems, it remains unclear whether simpler organisms also possess such capabilities. Here, we studied temporal information processing in *Schmidtea mediterranea*, a freshwater planarian flatworm with a primitive nervous system. Using long-term behavioral imaging and oscillatory ultraviolet (UV) light stimulations with rhythms matching the timescale of the animal’s short-term memory (∼minutes), we observed that planarians initially ignored rhythmic oscillations in UV intensity but eventually began tracking them after several cycles, demonstrating sensory filtering. We identified two neuropeptides, knockdown of which eliminated the initial ignoring phase and led to immediate stimulus-tracking, suggesting that these neuropeptides mediate an active sensory gating mechanism preventing response to transient fluctuations in stimuli. Notably, when UV stimulation was coupled with synchronous visible light oscillations, the planarians tracked the combined signals immediately, indicating that coherence across sensory modalities can override the initial gating. Our findings demonstrate that even simple nervous systems can filter temporal information and that this mechanism is mediated by neuropeptides. Unlike classical fast-acting small-molecule neurotransmitters, neuropeptides provide a slower, sustained, and global form of modulation that allows for more sophisticated control of sensory processing.

**Significance statement:** We show that simple nervous systems can use specific neuropeptides to achieve sensory filtering, a behavior previously thought to require complex brain architecture. This neuropeptide-mediated sensory gating mechanism reveals a fundamental role for neuropeptides in temporal information processing, offering insights into the mechanistic and evolutionary origins of attention-like behaviors.

## Introduction

Animals inhabit dynamic environments where they continuously encounter sensory inputs of various modalities and timescales, many of which may be irrelevant to their immediate needs. To navigate these complex conditions, it is beneficial to selectively respond to pertinent signals while filtering out extraneous information (1–3). This filtering also applies to temporal patterns, allowing animals to distinguish transient and persistent stimuli (4). Extensive studies in humans and other mammals have revealed complex neural circuits for sensory filtering (5–7). In invertebrates, such processes have been characterized in insects like *Drosophila*, which also rely on advanced brain structures such as mushroom bodies for sensory filtering (8–10). These observations have led to the notion that sensory filtering involves intricate neural circuits and dynamics (6, 11), though the underlying molecular mediators remain largely unknown. This raises an important question: Is sensory filtering exclusive to animals with intricate brains, or can it also arise in simpler organisms with rudimentary neural structures? Addressing this question may uncover core mechanisms of this important neural function and shed light on its evolutionary origins.

Here, we studied the freshwater planarian *Schmidtea mediterranea*. The simplicity of its nervous system is such that whether it is a true brain or primitive cephalic ganglia is still contested. Nevertheless, it consists of canonical neural cell types expressing conserved neurotransmitters and receptors (12, 13), and drives basic behaviors such as phototaxis, thigmotaxis, and chemotaxis (14–16). Recently, we found that planarians have short-term memory lasting a few minutes (17), suggesting that they may process temporal information.

To explore sensory filtering, we examined how planarians respond to oscillatory ultraviolet (UV) light stimulations with minute-scale rhythms, matching the timescale of their memory. Surprisingly, we found that planarians initially ignored rhythmic oscillations but eventually tracked them, while continuing to ignore irregular oscillations. Using RNA interference (RNAi) to perturb key components of neural communications, we identified two specific neuropeptides essential for the initial ignoring. Notably, pairing UV stimulus with concurrently oscillating visible light, planarians followed the rhythm immediately, indicating that coherent multisensory inputs can override the default filtering behavior. Overall, these results suggest that even a simple system can filter sensory information, which is governed by an active gating mechanism involving neuropeptides. This function allows animals to delay tracking of stimulus rhythms only after confirming their persistence.

## Results

### Long-term imaging reveals delayed tracking of rhythmic signals

To investigate how planarians process temporal information, we employed a long-term imaging platform described in our prior work (17). This setup exposes planarians to controlled stimuli over extended durations to precisely quantify their behavioral responses across many individuals. We subjected planarians to 30-minute trials of sinusoidal UV stimuli, with periods ranging between 2-4 minutes, as the planarian’s short-term memory peaks at ∼3 min (17). We chose sinusoidal waves to avoid discontinuities in the time derivative, which elicit strong aversive response in planarians (17, 18). If planarians responded solely to the current stimulation, we would expect their behavioral activity to track the UV oscillations, peaking in phase with the stimulus (**Fig. 1A**).

**Figure 1.**
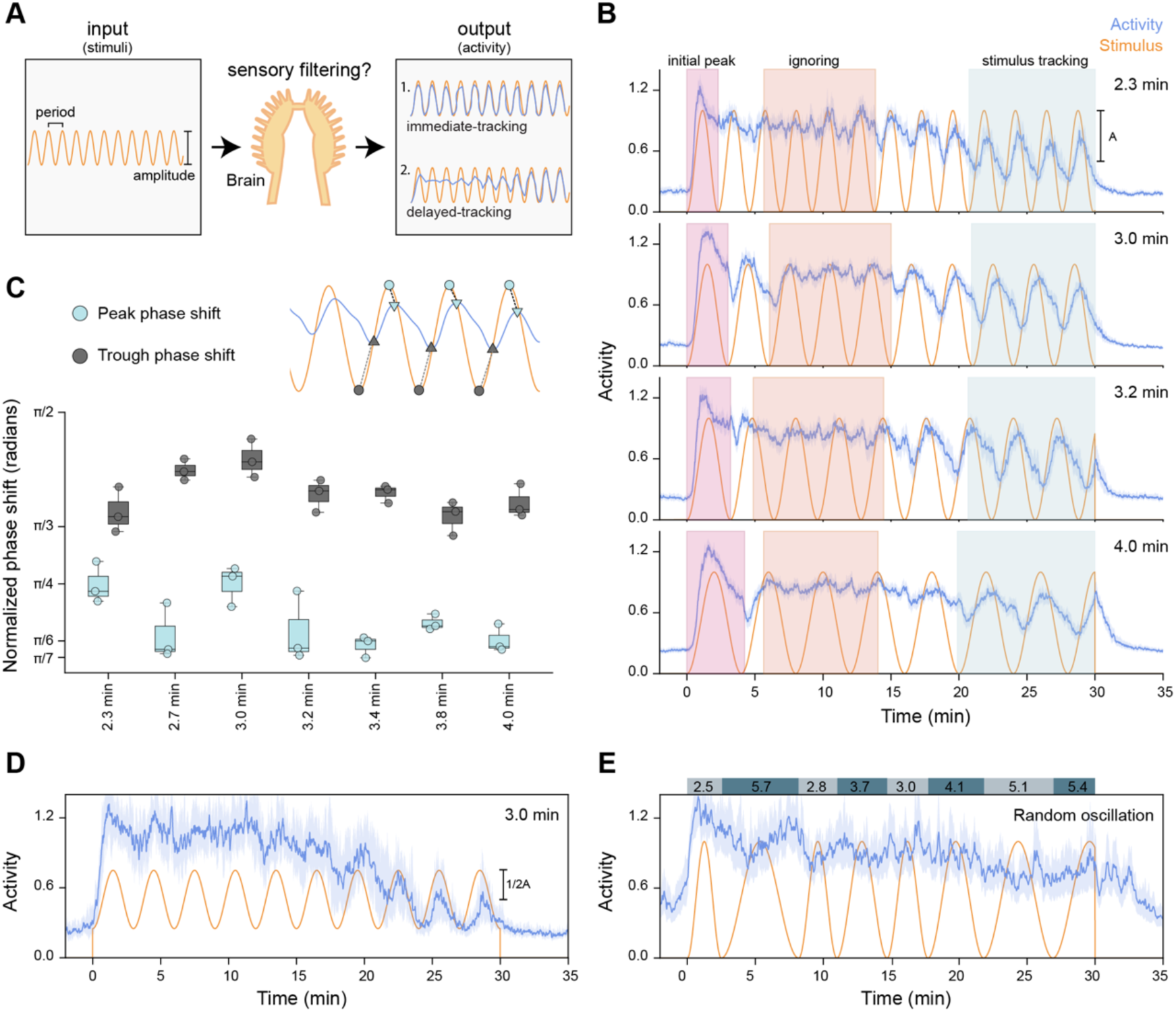
Planarians exhibit delayed stimulus-tracking in response to UV sine waves. A. Schematic of the behavioral measurement to study sensory filtering in planarians. Two potential outcomes are illustrated: immediate or delayed stimulus-tracking. B. Behavioral activity of planarians exposed to 30-min UV sine waves with four different periodicities. Highlighted sections: initial peak, ignoring phase, and stimulus-tracking. Blue lines: median activity; orange lines: stimulus profile. C. Top: diagram depicting the phase shift calculation. Bottom: phase shift for peaks and troughs during the last three cycles vs. stimulus periodicities. D. Activity under UV sine waves at half amplitude, showing a longer ignoring phase. The mean intensity of the stimulation is adjusted to match other conditions. E. Exposed to UV cycles with randomly generated periodicities results in no stimulus-tracking. **Statistics**: In (B, D-E), shaded regions: 95% confidence interval (CI); orange lines: stimulus profile. In (C), dots represent averages of the last three peaks from combined trials and experiments. The box-and-whisker plot shows the distribution of normalized phase shifts: boxes, interquartile range (IQR); bar, median; whiskers, 1.5 × IQR. Sample sizes: in (B), 2.3-min period: 2 batches/135 trials; 2.7-min: 3 batches/104 trials; 3.0-min: 7 batches/388 trials; 3.2-min: 3 batches/177 trials; 3.4-min: 2 batches/237 trials; 3.8-min: 1 batch/50 trials; 4.0-min: 8 batches/374 trails. In (D), 3 batches/102 trails; and in (E), 3 batches/60 trails. Each animal is exposed to a maximum of 10 trials, with each trial separated by a two-hour interval.

Upon UV exposure, planarians exhibited an immediate peak in behavioral activity, measured by a scalar metric that quantifies high-dimensional behavioral output (17) (**Fig. 1B**). This initial peak did not represent stimulus-tracking, as a similar response occurred upon exposure to constant UV **(Fig. S1A)**. Following the initial reaction, activity levels became relatively constant, indicating that the animals ignored the oscillations in stimulation strength. Surprisingly, after five to six cycles, they started tracking the UV oscillations, displaying clear peaks and troughs with a phase lag relative to the stimulus. This phase lag demonstrates that planarians were not merely reacting to immediate stimulation but were filtering temporal information based on its history. We quantified the phase lags by calculating the time difference between stimulus and activity peaks or troughs, normalized by the UV sine wave’s periodicity (**Fig. 1C**). The lags were consistently more pronounced in the troughs, revealing an asymmetry in the intrinsic neural processing delay.

To understand which behaviors contributed to these activity patterns, we used a Hidden Markov Model (HMM) to decompose behaviors into distinct movement types (17). This analysis revealed that roaming and nodding behaviors accounted for the observed activity oscillations, each showing a temporal profile that closely matched overall activity **(Fig. S1B)**. To determine whether the delayed stimulus-tracking resulted from a subset of “responder” animals gradually synching, we performed principal component analysis (PCA) on individual worms from multiple trials **(Fig. S1C)**. The analysis showed no distinct subpopulations, indicating that delayed tracking was a behavior consistent across individuals. Additionally, when we aggregated data based on trials, we found no clear evidence of long-term learning or memory across trials **(Fig. S1D)**.

Reducing oscillation amplitude, while keeping the mean intensity constant, decreases the animals’ ability to track stimulations (**Fig. 1D; Fig. S1E)**. At half amplitude, they began tracking the stimulus only after ∼8 cycles; at a quarter amplitude, the tracking was lost. Finally, when exposed to UV oscillations with changing periods, planarians exhibited high activity with no stimulus-tracking (**Fig. 1E**), suggesting that regular rhythmic input is necessary to induce tracking.

These findings suggest that planarians ignore transient oscillations before committing to a sustained tracking response to consistent and persistent stimuli. The observed initial delay and phase lag in tracking demonstrate a sophisticated temporal processing behavior of this simple nervous system.

### Neuropeptides mediate sensory filtering of oscillating signals

We hypothesized that the delayed tracking could result from either active gating or gradual adaptation/learning. If gating were the mechanism, disrupting its molecular mediators should prompt immediate tracking, whereas if adaption were involved, disruption should impair or prevent tracking (**Fig. 2A**). To test these possibilities, we used RNAi to knock down major neurotransmitters. Disrupting monoamine neurotransmitters did not significantly alter the tracking behavior. For example, knockdown of tyrosine hydroxylase (*th*), which inhibits dopamine synthesis (19), did not abolish the ignoring phase though it dampened the amplitude of the activity oscillations **(Fig. S2A)**. Knockdown of choline acetyltransferase (*chat*), which blocks acetylcholine synthesis (20), shortened the delay and exaggerated the response amplitude, consistent with our previous findings that acetylcholine is a major inhibitory neuromodulator in planarians (17), but it did not fully eliminate the ignoring behavior (**Fig. 2B**).

**Figure 2.**
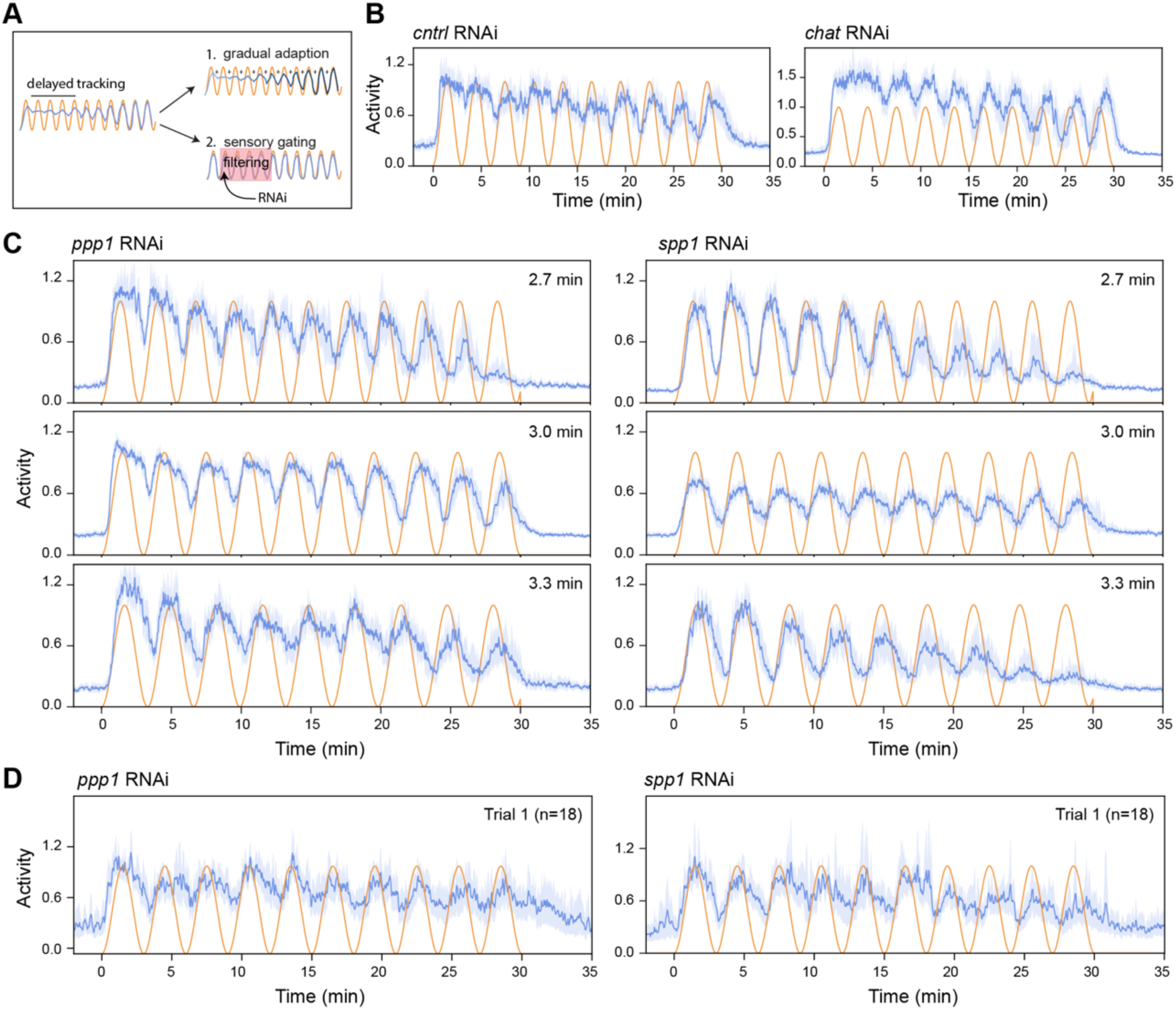
Neuropeptides regulate temporal sensory filtering. A. Schematic illustrating two potential mechanisms for the delayed stimulus-tracking that can be distinguished through RNAi experiments: (1) first gradual adaptation, or (2) active gating. B. Activity of control (left) and *chat* RNAi (right) planarians under UV sine wave stimulation with a 3-min period. Blue lines: median activity; orange lines: stimulus profile. C. Activity of *ppp1* (left) and *spp1* (right) RNAi planarians exposed to sine waves with periods of 2.7, 3.0, and 3.3 min. D. Activity of *ppp1* (left) and *spp1* (right) RNAi planarians during the first trial of UV sine wave stimulation with a 3-min period. **Statistics**: Shaded regions: 95% CI. Sample sizes: in (A), 2 batches/62 trials for control RNAi; 1 batch/70 trials for *chat* RNAi; in (B), for *ppp1* RNAi, 1 batch/50 trials for 2.7-min period; 4 batches/169 trials for 3.0-min period; 1 batch/54 trails for 3.3-min period; for s*pp1* RNAi, 1 batch/52 trails for 2.7-min period; 3 batches/98 trails for 3.0-min period; 1 batch/57 trials for 3.3-min period.

Given the essential roles of neuropeptides in regulating UV responses and short-term memory in planarians (17), we targeted several abundant neuropeptides, including *eye53*, *1020HH*, *spp-1*, and *ppp-1*, expressed in distinct cell types throughout the planarian brain (21–23). Strikingly, knockdown of *ppp-1* or *spp-1* caused immediate and sustained stimulus-tracking (**Fig. 2C**). Even during the first trial, *ppp-1* and *spp-1* RNAi animals followed the oscillation without any initial delay (**Fig. 2D**). These results suggest that the ignoring behavior is due to active sensory gating through these two specific neuropeptides. It is worth noting that *ppp-1* and *spp-1* knockdowns did not alter the temporal profiles of responses to short UV pulses **(Fig. S2B)** or constant UV exposure **(Fig. S2C)**, indicating that their role is specific to processing complex temporal information rather than general UV sensitivity.

Together, our results demonstrate that specific neuropeptides modulate sensory gating, delaying the tracking of oscillatory inputs. Given that neuropeptides are the largest and most diverse class of signaling molecules and are evolutionarily ancient (24, 25), their function in this primitive nervous system might represent a fundamental mechanism for modulating animal behavior in dynamic environments.

### Coherent multisensory inputs override gating

In natural environments, sensory inputs from multiple modalities coexist, which require integration for appropriate behavioral responses. For planarians, UV light and visible light are typically concurrent in their natural habitat of shallow water, presenting a natural scenario where dual inputs must be processed together. *S. mediterranea* possesses distinct ocular and extraocular photoreceptors that detect visible and UV light, respectively (15). This separation allowed us to independently stimulate these two sensory modalities to investigate how planarians integrate inputs sensed differently.

When exposed to oscillatory visible light (520 nm), planarians showed no stimulus tracking (**Fig. 3A**), suggesting that visible light alone is insufficient to evoke a tracking response, despite its ability to induce strong negative phototaxis (14). To our surprise, when exposed to simultaneous oscillations of UV and visible light, planarians tracked the combined stimuli without delay, similar to the response observed in *ppp-1* and *spp-1* knockdown conditions (**Fig. 3B**). This indicates that coherent multisensory inputs can override the default sensory gating.

**Figure 3.**
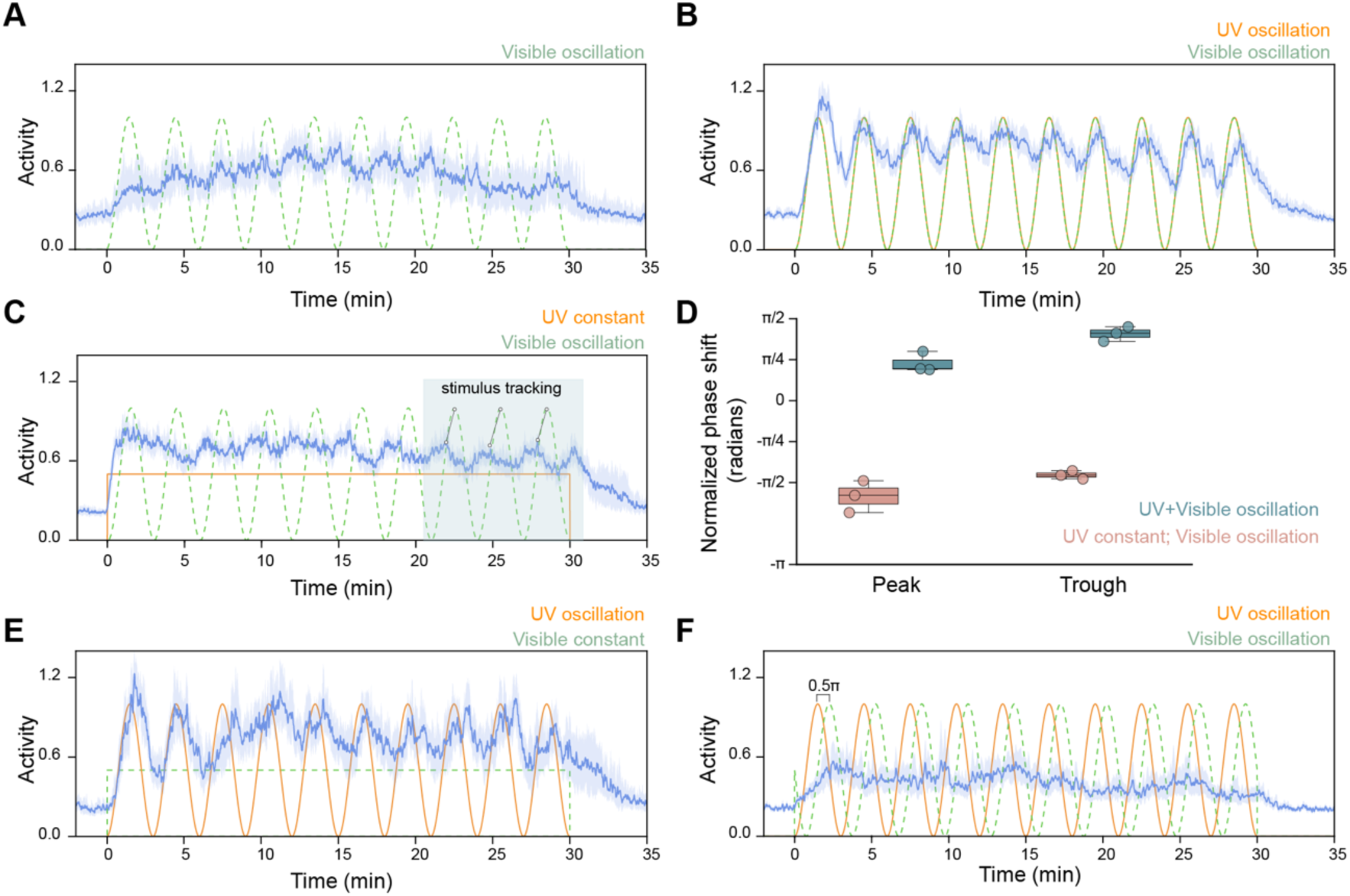
Coherent multisensory inputs override sensory filtering. A. Activity under visible light sine wave with a 3-min period. Blue line: median activity; green dashed line: visible light stimulus profile. B. Activity in response to coherent UV and visible light stimulations, both applied as sine waves with a 3-min period. Orange line: UV stimulus profile. C. Activity in response to a constant UV stimulus, set to the mean intensity of the sine wave stimulus, combined with a visible light sine wave with a 3-min period. Highlighted: stimulus-tracking in the last three cycles. This condition induces a phase lead in tracking behavior, indicated by the gray line. D. Quantification showing the flip of phase shifts under conditions shown in (B, blue) vs. (C, red). Box plot displays normalized phase shifts for the last three peaks and troughs. E. Activity response to a continuous visible light stimulation combined with a UV sine wave with a 3-min period. F. Activity response to a UV and visible light sine waves with a 0.5π phase shift. **Statistics**: In (A-C, E-F), shaded regions: 95% CI. In (D), dots represent the average of the last three peaks from combined trials and experiments. The box-and-whisker plot shows the distribution of normalized phase shifts as in Fig. 1C. Sample sizes: 2 batches/79 trials (A); 2 batches/92 trials (B); 2 batches/108 trials (C); 2 batches/68 trials (E); 2 batches/77 trials (F).

We further tested responses to constant UV light with oscillatory visible light and found that planarians tracked visible light oscillations (**Fig. 3C**), but with a phase lead, as though predicting upcoming changes (**Fig. 3D**). In contrast, combining oscillatory UV and constant visible light replicated the delayed tracking observed for UV alone (**Fig. 3E**). Lastly, coupling UV and visible light oscillations with a phase shift of 0.5π, eliminated stimulus-tracking (**Fig. 3F**), highlighting the importance of coherence between sensory modalities for overriding filtering and guiding different behavioral outcomes.

These findings demonstrate that the sensory filtering mechanism in planarians is not limited to UV stimulation alone but can be modulated or overridden by inputs from other sensory modalities, highlighting the relevance of sensory filtering in differentiating various types of multimodal signals. These insights may also help understand basic principles of multisensory integration.

## Discussion

Our study demonstrates an unexpected simplicity underlying the seemingly complex neural function of sensory filtering: it exists within a simple nervous system and can be disrupted by knocking down individual neuropeptides or overridden by coherent multisensory inputs. Since neuropeptides are abundantly used in organisms (24–26), our results may have broad implications.

We propose two potential mechanisms through which neuropeptides mediate temporal sensory filtering. First, unlike classical neuromodulators, which act rapidly at synapses, neuropeptides can diffuse over long distances and persist in the extracellular space for extended time (27, 28). This widespread diffusion could synchronize neural activity across space and/or time, a process thought to be necessary for selective filtering or attention-like behaviors (6). Alternatively, the observed gating might result from neural circuits dedicated to integrating sensory inputs that are modulated by neuropeptides. The relatively slow action of neuropeptides could naturally provide the nervous system with a memory on the minutes timescale necessary for sensory filtering (9). The inertia to changes in the neural state could prevent the organism from overreacting to transient stimuli, allowing it to focus on persistent and relevant environmental cues. In either case, our work highlights the function of neuropeptides in modulating behaviors that need to evoke long-lasting temporal neural states (29).

What determines the timescale of filtering? It is plausible that the timescale reflects the ecological significant cues in planarians’ natural environment. Planarians lack vision, and their light perception primarily relies on encoding light intensity and wavelength (18). Abrupt changes in illumination may signal immediate threats, such as the sudden appearance of predators or habitat boundaries, prompting instant response; even gradual changes in light intensity, analogous to “looming stimuli” in mouse vision test (30), can indicate a predator approaching or retreating thereby requiring a consistently active response. Persistent rhythmic fluctuations, on the other hand, may convey different information, such as variable light patterns caused by water movement, and demand a different response. By filtering out initial fluctuations in sine waves, planarians can allocate their limited neural resources to different aspects of the environmental cues based on the persistence of temporal patterns.

Alternatively, our findings may reveal neural processes that are not directly linked to specific environmental cues but are intrinsic to the operating mechanisms. In small nervous systems, neuropeptides can have disproportionately large effects, potentially inducing slow neural dynamics that influences global brain states. These small peptides (<40 amino acids), packaged in dense core vesicles, are released in response to depolarization at various sites along the neuron (28, 31). Their sizes and structural features contribute to a longer half-life on the scale of hundreds of seconds (28, 32, 33), and the lack of specific reuptake mechanisms means that neuropeptides remain in the extracellular space longer (33). Assuming simple diffusion, we estimate that neuropeptides can travel extracellularly and influence neural activity over distances of >500 μm. While this distance is relatively small in large, complex brains, it is almost the entire dimension of the planarian brain, implicating that peptides can induce global states across the brain. Consistently, in animals of similar or even smaller sizes, such as *Caenorhabditis elegans*, the neuropeptide connectome illustrates how neuropeptides can bridge otherwise disconnected neural circuits, forming a dense and decentralized signaling network (34). Indeed, in *C. elegans*, most active neurons in the brain participate in continuous coordinated neural activity fluctuations on the scale of ∼100 s to represent various behaviors, including sensory-driven action selection (35, 36).

To determine whether neuropeptide-mediated filtering arises from modulation of neural synchrony, specific circuit dynamics, or a combination of both requires direct measurements of neural activity. Although technical limitations currently preclude *in vivo* calcium imaging in planarians (37), our behavioral paradigm using UV sine waves to probe temporal filtering can be applied to other models with advanced genetic and imaging tools (38, 39). Ultimately, understanding how neuropeptide-mediated processes contribute to sensory processing across different species and contexts may inform general principles of sensory filtering and its evolution.

## Materials and Methods

### Animal care and maintenance

Asexual *S. mediterranea* were maintained in the dark at 20 °C in water containing 0.5 g/L Instant Ocean Sea Salts and 0.1 g/L sodium bicarbonate. Behavior experiments used planarians of ∼4 mm in length. Animals were fed every 4-7 days and starved a minimum of 4 days before behavioral recording.

### Imaging setup

The imaging setup, detailed in ref. (17), illuminated animals with an IR light (850 nm) and recorded at 2 frames per second using a Raspberry Pi NoIR camera, ensuring minimal interference with their natural behavior. UV stimuli (365 nm) were delivered by a custom-built ring of 36 LEDs mounted above the camera to achieve uniform illumination across the dish, and controlled by an Arduino Uno for precise timing and intensity modulation. Visible light (520 nm) was similarly controlled and delivered using the Adafruit NeoPixel RGB LEDs (model 1586).

In all stimulation experiments, we maintained a two-hour period of unstimulated, dark time between repetitions of the protocols to prevent any influence between trials or cumulative effects on behavior. A total of 24 hours of data was collected for each experiment, corresponding to 10 trials.

### RNAi

Gene knockdowns were performed by feeding animals double-stranded RNA (dsRNA). The dsRNA was synthesized following the standard protocol(21) and fed to the planarians via a liver homogenate at a concentration of ∼100 ng/µL. Clones for dsRNA synthesis were created using oligonucleotide primers reported in ref. (17) and cloned into the vector pJC53.2 (Addgene plasmid ID: 26536) (21). Plasmids containing neuropeptide sequences were from ref. (21).

For the RNAi experiments, animals were fed dsRNA 5-7 times at 4-5 days intervals. For the controls, animals were fed dsRNA matching the ccdB and camR insert of pJC53.2 in parallel. All animals were then starved for 4 days prior to decapitation, after which the tails were allowed to regenerate and were imaged after regeneration completed (at 10-15 days post-amputation).

### Behavioral activity quantification

The methodology for quantifying planarian behavior was adapted from ref. (17). Behavioral data were processed and analyzed to determine patterns of activity in response to UV and visible light stimuli. To assess statistical significance, a bootstrap resampling method was employed with 1,000 bootstrap samples, allowing for reliable estimation of confidence interval (CI) around the median activity values.

### Data and code availability

Code for image segmentation is available at github.com/samuelbray32/planameterization (https://doi.org/10.5281/zenodo.12697208). Code for data analysis and visualization is available at github.com/lwyss/timescales_behavior. We acknowledge the use of ChatGPT for assistance in simplifying and annotating code used for plotting the activity data.

## Acknowledgements

We thank Liqun Luo, Lauren O’Connell, Chris Lowe, Prateek Kalakuntla, Jesse Gibson, and other Wang lab members for valuable discussions. LSW and SRB acknowledge the support from a NIH cellular, Biochemical, and Molecular Sciences (CMB) training grant (T32GM007276) and NSF GFRP fellowships. This work is supported by an NIH grant (1R35GM138061) and the Neuro-omics project of Wu Tsai Big Ideas in Neuroscience program.

## Supplementary Figures

**Figure S1.**
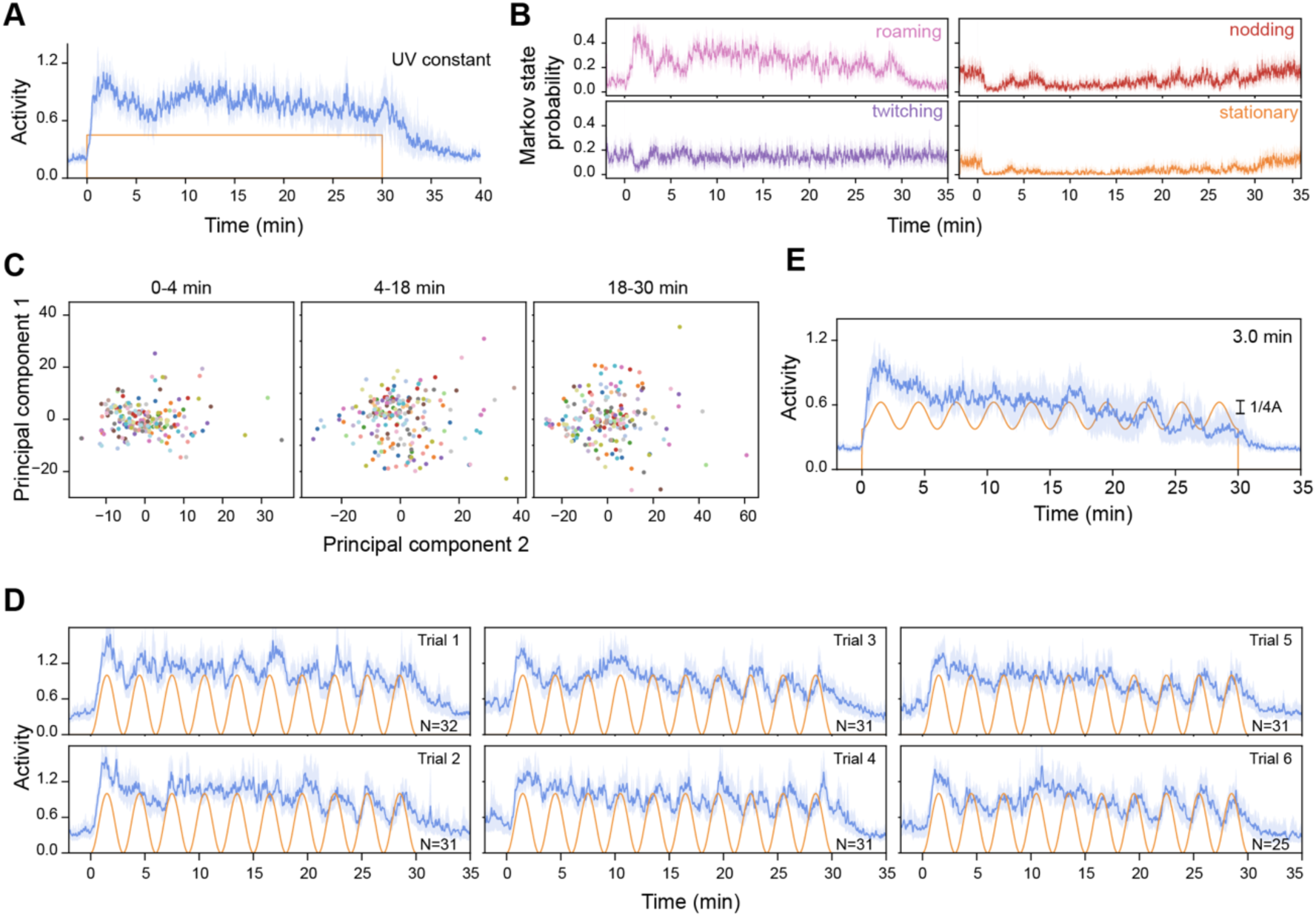
Additional characterization of stimulus-tracking behavior. A. Planarian activity under continuous UV stimulation for 30 min, matched in average intensity to the sine wave stimulations. Blue line: median activity; orange line: stimulus profile. B. Observed probability of HMM states during UV sine wave stimulation with a 3-min period. Oscillations (stimulus-tracking) are clear in roaming and nodding states. C. PCA of individual animal activity, with each animal represented by a unique color. Data in (B, C, D) is from the 3-min period experiment shown in Fig. 1B. D. Activity traces grouped by trial number for the first six trials, showing consistent activity patterns with no apparent trial-to-trial differences or long-term learning effects. E. Activity profile under UV sine wave at quarter amplitude (a 3-minute period). The mean intensity of the stimulation is adjusted to match other conditions. **Statistics**: In (A, E, D), shaded regions: 95% CI. Sample sizes: 1 batch/48 trails (A); 3 batches/92 trials (E). In (B), the explained variance by the first two PCs: initial peak (0–4 min): PC1 = 26.62%; PC2 = 16.63%; ignoring phase (4–18 min): PC1 = 16.32%; PC2 = 10.65%; stimulus-tracking (18–30 min): PC1 = 26.40%; PC2 = 11.68%.

**Figure S2.**
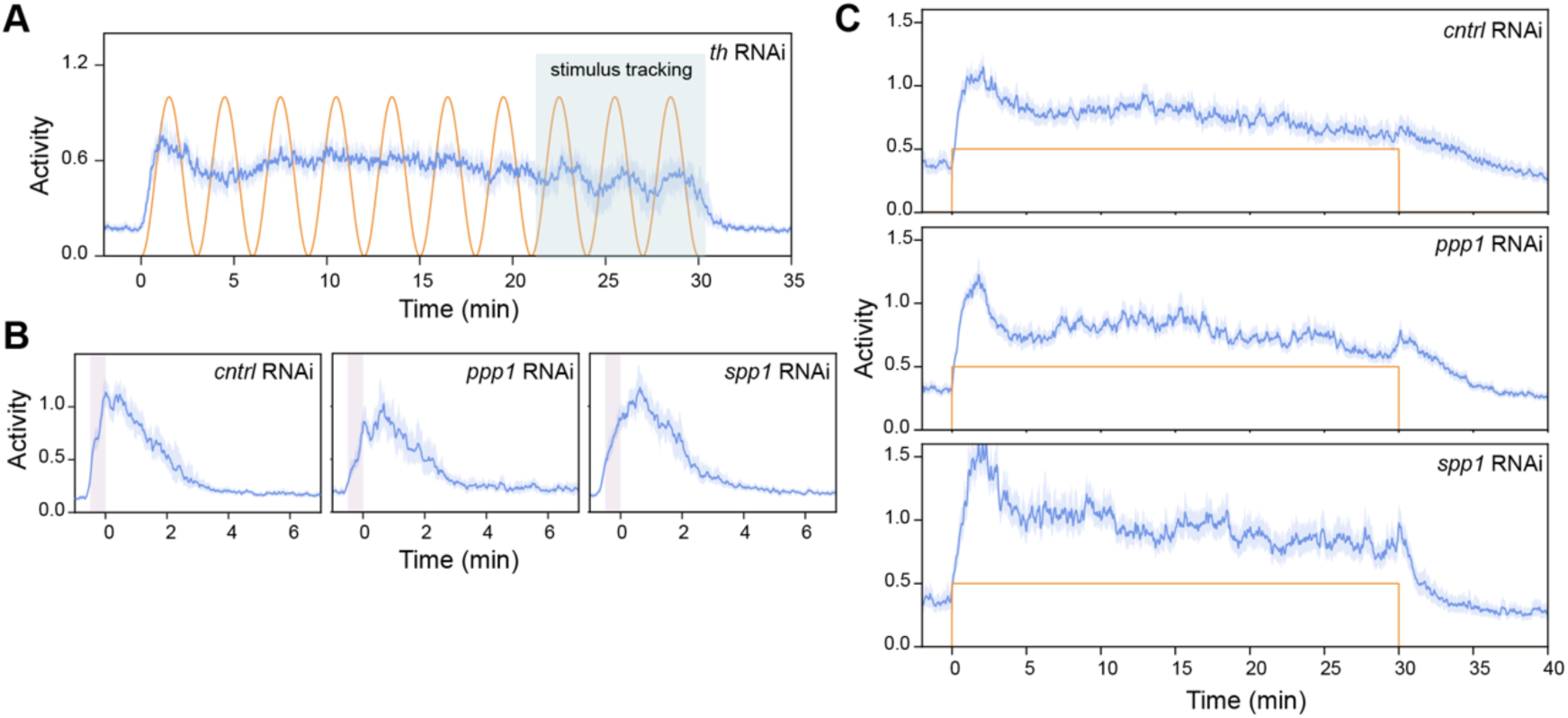
Additional characterization of UV responses in RNAi-treated animals. A. Activity of *th* RNAi planarians subjected to UV sine wave stimulation with a 3-min period. Highlighted: stimulus-tracking in the last three cycles. Blue line: median activity; orange line: stimulus profile. B. Activity of RNAi-treated animals in response to a 30-second UV pulse ending at time 0, which show no significant difference across conditions. C. Activity of RNAi-treated animals under continuous 30-min UV exposure, with intensity matching the mean intensity of sine wave stimulations. **Statistics**: Shaded regions: 95% CI. Sample size: in (A), 2 batches/87 trials; in (B), 2 batches/108 trials for control RNAi; 1 batch/60 trails for *ppp1* RNAi; 1 batch/96 trials for *spp1* RNAi; in (C), 2 batches/142 trials for control RNAi; 3 batches/136 trails for *ppp1* RNAi; 2 batches/313 trials for *spp1* RNAi.

